# Alpha-Synuclein Fibril Structures Cluster into Distinct Classes

**DOI:** 10.1101/2025.04.30.651534

**Authors:** Moses H. Milchberg, Owen A. Warmuth, Collin G. Borcik, Dhruva D. Dhavale, Elizabeth R. Wright, Paul T. Kotzbauer, Chad M. Rienstra

## Abstract

The accumulation of Alpha-synuclein (Asyn) fibrils is the defining pathologic feature in Parkinson Disease (PD), Lewy Body Dementia (LBD), and Multiple System Atrophy (MSA). As such, the process of Asyn fibril formation has been an important research area and fibrils themselves have become attractive targets for disease diagnosis and therapeutic intervention. Due to the presence of mixed populations of fibrillar proteins associated with neurodegenerative diseases in brain tissue, high-resolution structures of Asyn fibrils are essential for the design of high-specificity imaging and therapeutic agents. Approximately one hundred high-resolution solid-state NMR (SSNMR) spectroscopy and cryo-electron microscopy (cryo-EM) structures of Asyn fibrils have been deposited to the Protein Databank (PDB); intriguingly there is significant polymorphism among them. Understanding the molecular makeup and characteristic features of each structural polymorph can determine conserved structural motifs which can be used as templates to design ligands with high specificity for clinical use. Utilizing standard alignment tools and density-based clustering approaches, we objectively classify fibril structures by tertiary structure type. We find that 81% of the structures cluster into two polymorph classes. Within each class, additional subtle variations are observed which position sidechains in specific, conserved orientations, well poised as druggable targets. Furthermore, we find that the conserved structural motifs associated with each class are found in all but one published Asyn fibril structure. We consider these classifications and conserved motifs in the context of disease-relevant fibril structures and offer a perspective on the utility of *in vitro* fibrils as substrates for drug development and models for disease pathogenesis.

## INTRODUCTION

Many neurodegenerative diseases, such as Parkinson disease (PD), Alzheimer disease (AD) and Huntington’s disease, are characterized by the misfolding and aggregation of proteins into amyloid fibrils, which deposit into extracellular intraneuronal plaques within the brain (1-3). One such protein, Alpha-synuclein (Asyn), natively exists as an intrinsically disordered monomer within the presynaptic termini of neurons and is involved in the organization and maintenance of synaptic vesicles under normal physiology (4,5). Asyn misfolds and forms fibrils as part of the etiology of PD, Lewy Body Dementia (LBD) (which includes Parkinson Disease Dementia (PDD) and Dementia with Lewy Bodies (DLB)), and Multiple System Atrophy (MSA), as well as other hereditary and sporadic disorders collectively known as synucleinopathies (6-9). The identification of autosomal dominant mutations in the gene (SNCA) encoding Asyn in hereditary forms of PD/LBD implicates Asyn in a central role in pathogenesis (10).

How physiological Asyn assembles into fibrils has been a high-interest area of study (11-15), with differences in formation kinetics and morphology observed between *in vitro* and *ex vivo* fibrils of distinct disease phenotypes (16,17). Targeting these fibrils within the brain offers an attractive diagnostic and therapeutic avenue for combating synucleinopathies. Historically, this approach has been unsuccessful due to the presence of other protein fibrils such as tau and amyloid-beta at high quantities in the brains of synucleinopathy patients (18,19). Ligands, such as Pittsburgh Compound B, have been successfully used as diagnostic positron emission tomography (PET) imaging agents in the case of amyloid-beta fibrils in AD, but have been shown to bind Asyn fibrils with weak affinity and specificity, preventing their use in the clinic as a potential diagnostic for synucleinopathies (20,21). The computational design of drugs with high specificity and affinity for Asyn fibrils has been shown to benefit from atomic level fibril structural information. Over the past 10 years, advancements in fibril structure determination methods specifically through solid-state NMR (SSNMR) spectroscopy and cryo-electron microscopy (cryo-EM), have expanded the < 4 Å resolution library of structures that are effective for computational approaches and have resulted in novel small molecule ligands with high specificity for Asyn fibrils (22-25).

Although the expanding number of available Asyn fibril structures has proved instrumental for rational drug design, it has also revealed that structural polymorphisms are highly sensitive to fibril growth conditions. Co-factors such as metals (26), ligands like heparin and lipids (27,28), and hereditary mutants (29,30) have resulted in a broad range of fibril structures. Additionally, differences in pH, buffer components and salt concentrations are also associated with the nucleation and formation of different conformations (31,32). Densities postulated to be these cofactors and ligands are commonly found within cryo-EM maps from both *in vitro* and *ex vivo* samples, though often the exact chemical makeup of these densities remains dubious (28,33,34).

While numerous structures have been determined over the past decade, their polymorphic nature has obfuscated determination of their disease relevance, utility as substrates for drug development and models for disease progression. *Ex vivo* structures from diseased patient tissue have revealed that while structural similarities are present, *in vitro* fibril preparations do not exactly capitulate the fold(s) found within diseased brains (33). There are several existing reviews of published Asyn fibril structures that have focused on on cryo-EM structures of *ex vivo* fibrils (35,36), or the analysis of conserved structural motifs (37-39). However, what was missing is an objective and statistical analysis of the entire PDB structural library of Asyn fibrils. This approach can provide a classification of the structural polymorphisms present in Asyn fibrils.

In this review, we focus on differences in fibril tertiary structure. We first identify a common fibril core present in each structure and align them. We then use principal component analysis (PCA) and density-based clustering to classify Asyn fibril protomers based on their PDB coordinates. This provided us with two major conformations of Asyn fibrils for further analysis. Within these major conformations, we identify conserved structural motifs and subclusters, placing them in the context of disease-relevant fibrils. Our classification indicates that certain *in vitro* fibrils are good models for studying the pathogenesis and progression of MSA and should be used as templates for drug development.

## ALPHA-SYNUCLEIN FIBRIL STRUCTURES AT A GLANCE

As of June 2024, 84 structures of Asyn fibrils have been published and deposited to the PDB. Among those, two were incubated with other proteins and 11 were bound to ligands (25,40-42). Here, we focus on the remaining 71, formed by *in vitro, ex vivo* and *ex vivo* seeded preparations (**Table S1**) (26-34,42-64). **Figure 1** provides a visualization of these preparations and details the proportion of structures prepared from these methods. The majority of the structures (∼65%) are from *in vitro* preparations, which includes five structures from sequence truncations or insertions (48,55,59), nine structures of point mutations (29-31,49,50,53,54,60), five structures with post-translational modifications (51,61,63) and 11 structures of fibrils formed in the presence of lipids or other cofactors (26-28). The second highest proportion of structures is *ex vivo* seeded – these include fibrils extracted from postmortem brain tissue or cerebral spinal fluid (CSF) which are then amplified *in vitro* (46,52,56-58). *Ex vivo* fibrils – those extracted from PD, PDD, DLB, MSA, and Juvenile Onset Synucleinopathy (JOS) patient tissue – represent the smallest proportion of fibril structures (33,34,59). There are ongoing questions whether seeding can faithfully template the *ex vivo* structures or introduce new primary nucleation events to generate entirely new structures (52,56-58).

**Figure 1:**
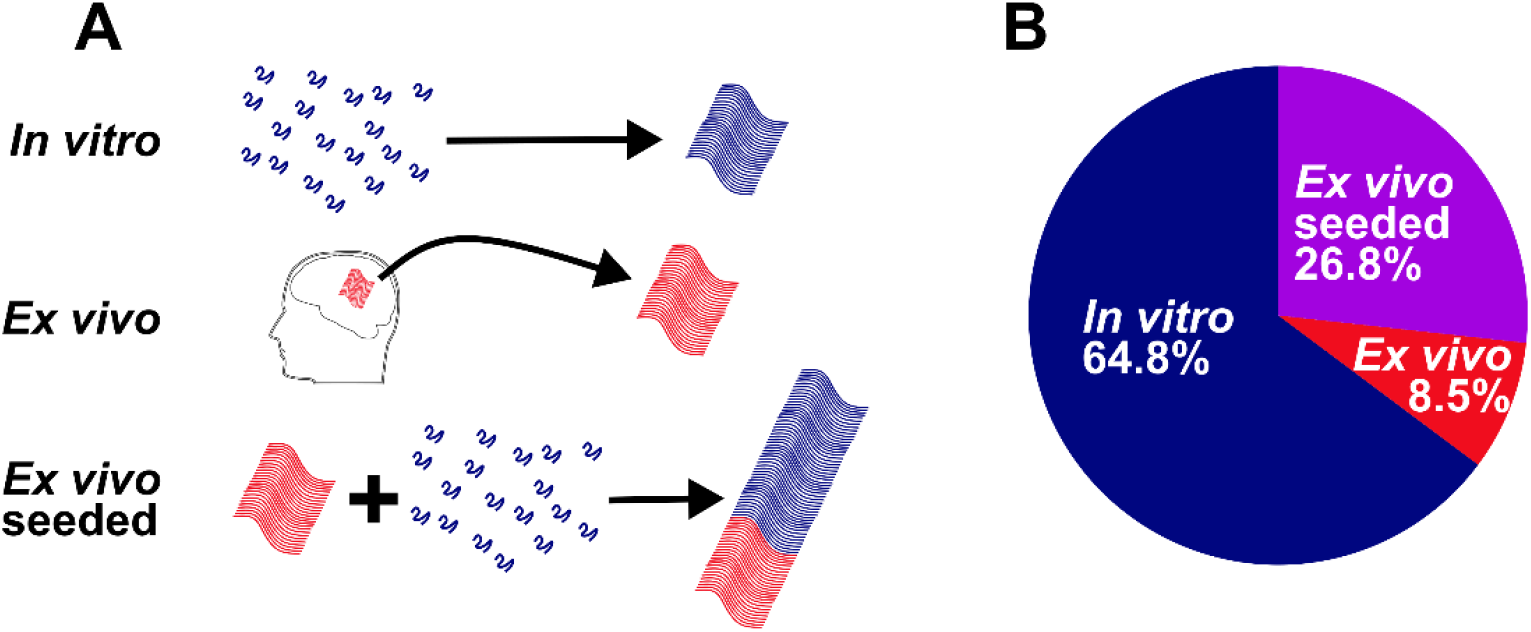
Preparations of Alpha-synuclein fibrils. (A) Schematic showing fibrils recombinantly expressed and formed *in vitro*, extracted from the brains of post-mortem patients (*ex vivo*) and amplified from CSF or brain tissue extract of patients (*ex vivo* seeded). (B) Out of the 71 apo structures published and deposited to the PDB, 46 are from *in vitro* preparations, 6 are extracted from *ex vivo* patient tissue, and 19 are seeded from *ex vivo* patient tissue or CSF.

Each structure has an ordered fibril core between residues 40-100 as well as dynamic and heterogenous termini. These structures are polymorphic, detailing differences in secondary, tertiary, and quaternary structure. For fibrils, secondary structure refers to the location of β-strands and turns, tertiary structure is the orientation of β-strands and turns within a protomer and is referred to as the fold of the fibril, and quaternary structure details the existence of multiple protofilaments and orientation of their interface.

### CLUSTERTING OF ALPHA-SYNUCLEIN FIBRIL STRUCTURES

We applied a quantitative and objective approach to our classification using the Bio3D package in R (65). We aligned the 75 unique protomers (four of the fibrils had asymmetric protofilaments) using MUltiple Sequence Comparison by Log-Expectation (MUSCLE) (66), among which 68 had a continuous fibril core spanning residues E46-A91 (**Figure S1)**. We used PCA to reduce the dimensionality (67-71) and Density-Based Spatial Clustering of Applications with Noise (DBSCAN) (72,73) to identify two distinct clusters, with Cluster 1 containing 28 protomers, Cluster 2 containing 27 protomers, and 13 remaining unclassified due to lack of proximity in PC space (**Figure 2A**). Both clusters of protomers were overlaid (**Figure 2B-C**), and a representative set of the unclassified protomers are shown side-by-side (**Figure 2D)**. Plots of their per-residue root-mean-square fluctuation (RMSF) are shown in **Figure S2** (74). Protomers which nearly overlap in PC space, such as 6L1U and 6L1T (pY39), have nearly identical folds yet have different filament-filament interactions between multiple protomers at the level of quaternary structure.

**Figure 2:**
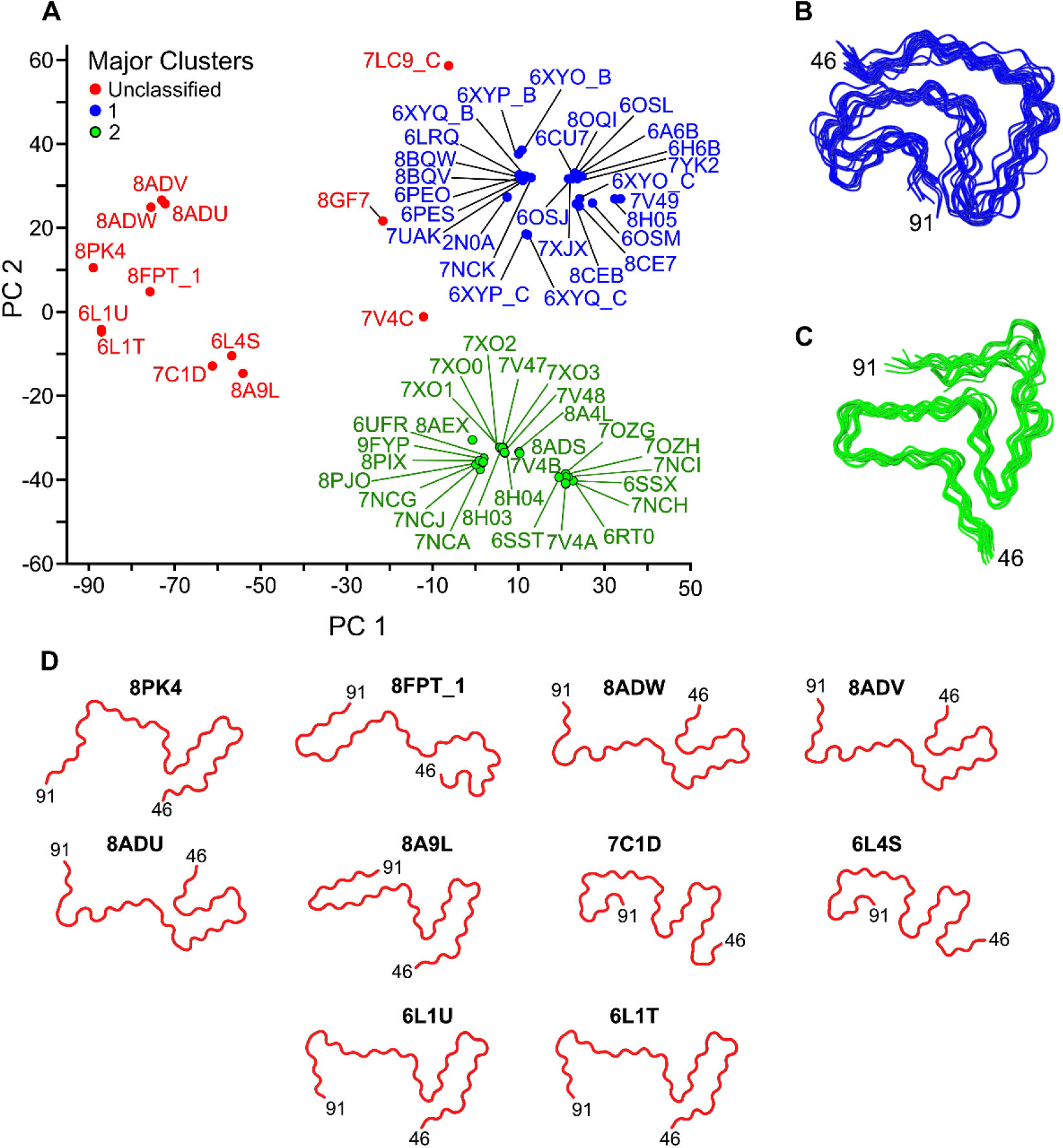
Objective clustering of Asyn fibril protomer structures. (A) PCA plot of the subset of PDB coordinates aligned from E46-A91. Each point represents the protomer of a single PDB coordinate file and is labeled with its four-character identifier and colored according to its DBSCAN cluster. Points colored in blue are part of Cluster 1, green are part of Cluster 2, and red are unclassified. (**Table S1**). Four-character identifiers with an underscore and letter following them denote the specific PDB chain ID, in cases where there are mutliple protofilaments that are asymmetric. 8FPT_1 dentoes the lowest energy protomer of 8FPT from the NMR structural bundle deposited to the PDB. Each PDB file within (B) Cluster 1 and (C) Cluster 2 were overlaid and aligned in ChimeraX, with average RMSFs of 2.31 Å and 1.99 Å shown in blue and green, respectively. (D) Ten of the 13 unclassified protomers shown side-by-side in a grid with residues 46 and 91 labeled. Similarities in structure explain some of their close proximity in PC space even though as a whole they do not form a Cluster.

### CLUSTER 1 STRUCTURES

Cluster 1 includes the first structure of full-length Asyn fibrils by SSNMR (2N0A), 16 additional *in vitro* fibril preparations, two *ex vivo* seeded PD structures, one *ex vivo* seeded MSA structure, six *ex vivo* MSA structures, and two *ex vivo* JOS structures. The aligned structures have an average RMSF of 2.31 Å (**Figure S2B**), illustrated in **Figure 3A** where the SSNMR structure is shown in blue. Additional differences are observed in the C-terminus of the core, with the RMSF increasing to at least 4.0 Å for each residue between A85-A91 (**Figure S2B)**. The overall fold contains the following four motifs which contribute to its stability: (1) Steric zippers forming between the G47-A53 and V74-Q79 βstrands, including sidechain interactions among residues A78-V49-A76, V74-A53, V77-A90, and T75-T92-V71 (**Figure 3C-D**); (2) an orthogonal Greek key forming among residues A69-V95, usually including the close proximity of the I88, A91 and F94 sidechains in a hydrophobic pocket (**Figure 3B**); (3) intermolecular salt-bridges forming between E46-K80 (**Figure 3F**) and (4) a glutamine ladder at Q79 (**Figure 3E)** forming between each filament along the fibril axis.

**Figure 3:**
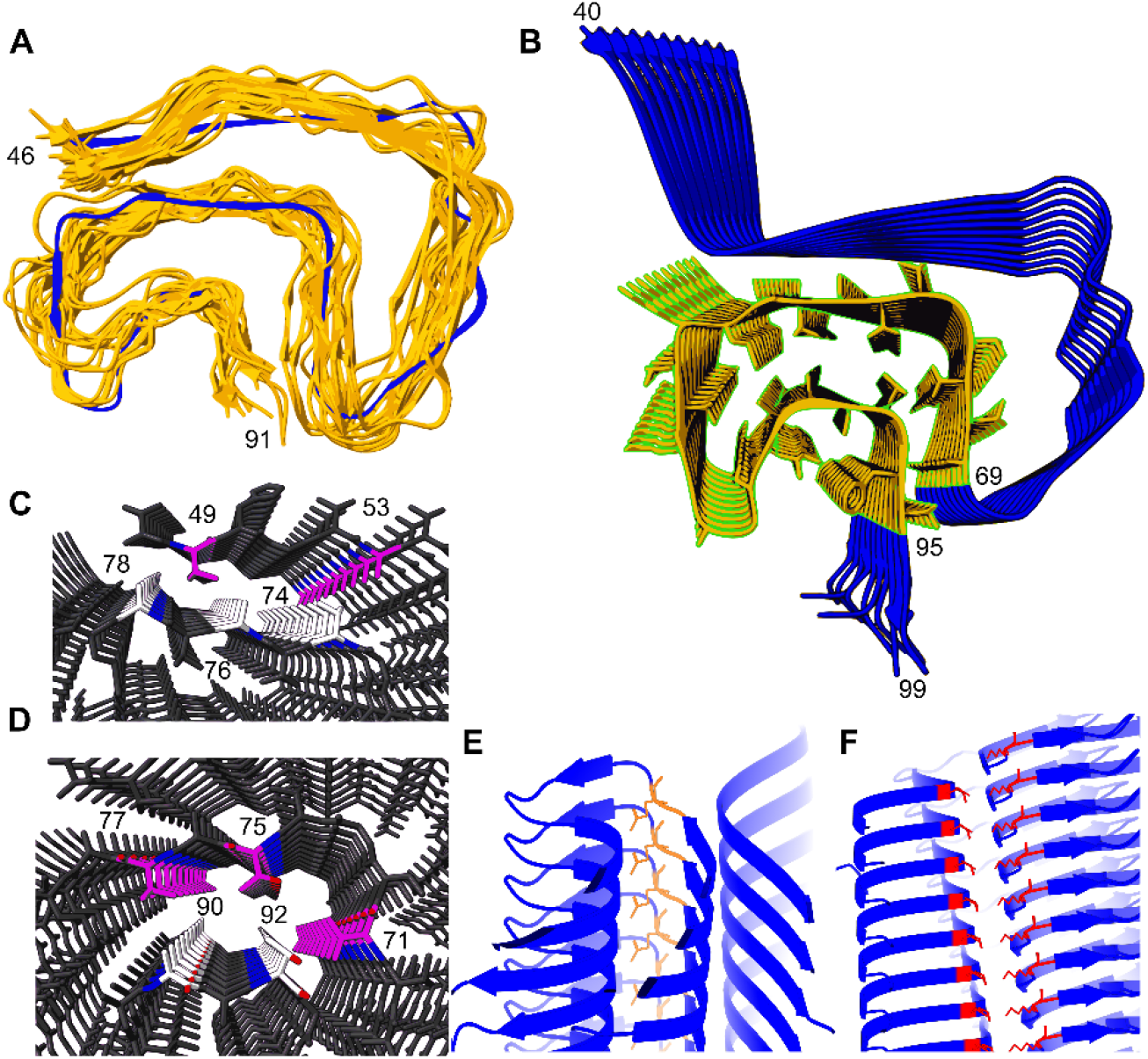
Characteristic features of Cluster 1 polymorphs. (A) Overlay of all Cluster 1 structures in orange with 2N0A in blue, aligned from E46-A91. (B) Truncated V40-Q99 peptide of 2N0A showing characteristic Greek key region between A69-V95, with a hydrophobic pocket featuring I88 and F94 sidechains in close proximity. Close up view of fibril detailing steric zippers forming between (C) A78-V49-A76, A53-V74 and (D) V77-A90-T75-T92-V71, with carbons on opposing beta strands colored in purple and white, nitrogen atoms colored in blue, and oxygen atoms colored in red. Side view of (E) glutamine ladder at Q79 (orange) and (F) E46-K80 intermolecular salt bridge (red).

Even though there is overall agreement within the protomers of Cluster 1, we sought to quantify the heterogeneity and identify its residue-specific determinants. To do this, we repeated PCA on the Cluster 1 structures. In the multiple sequence alignment **(Figure S1)**, all of the Cluster 1 protomers had continuous fibril cores spanning E46-F94, thus we used those residues for the alignment and PCA, which separated Cluster 1 into two subclasses, based on the differences in PC1 (**Figure 4A**). The average RMSF for Cluster 1 (E46-F94) is 2.59 Å, with subclass differences substantiated by the increase in per-residue RMSF to > 4.0 Å between A85-F94 (**Figure 4B**).

**Figure 4:**
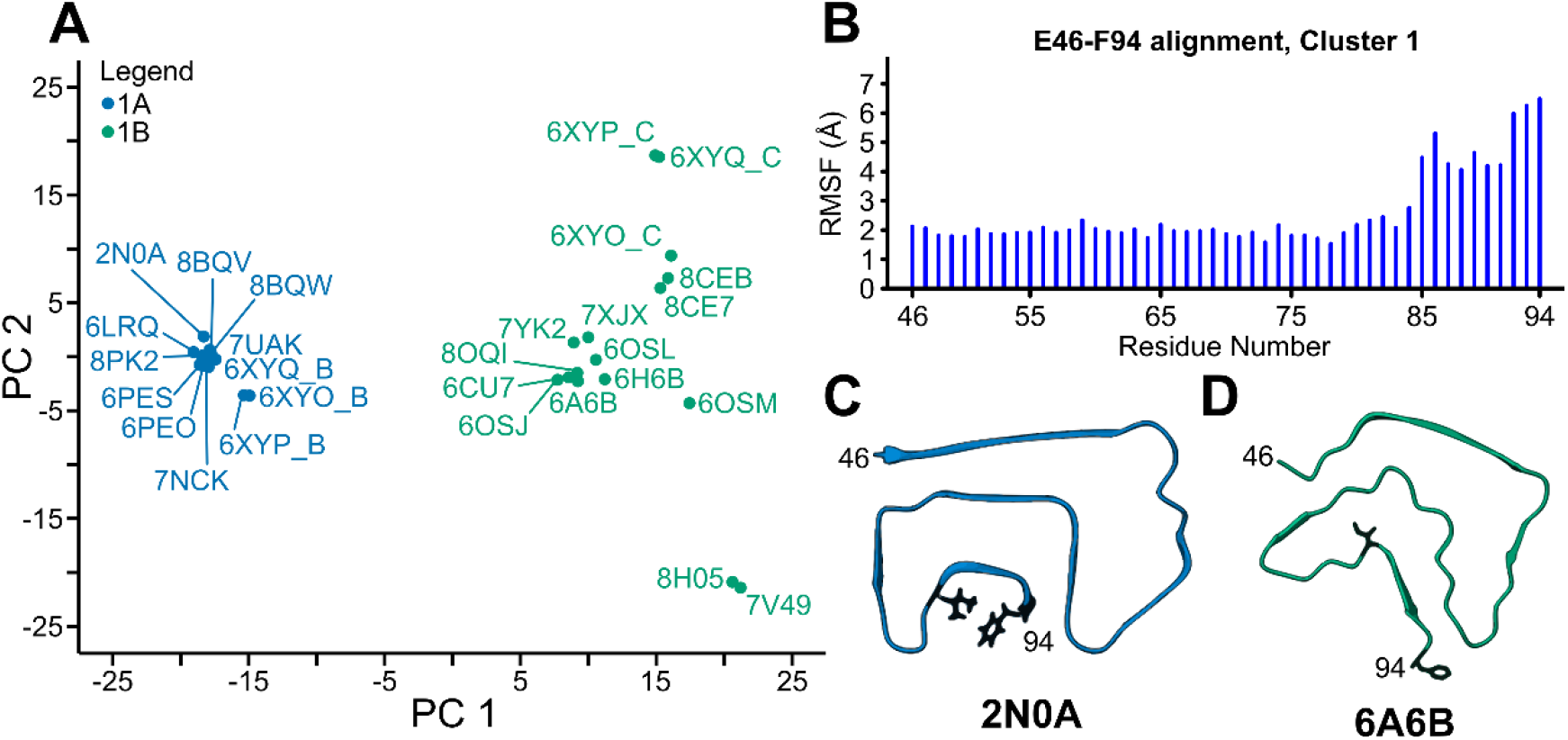
Identification of subclasses within Asyn fibril Cluster 1 protomers. (A) PCA plot (PC 2 vs PC 1) of all protomers within Cluster 1, aligned from E46-F94. Each point represents the protomer of single PDB coordinate file and is labeled with its 4-character identifier, with points corresponding to Cluster 1A (negative PC 1 value) colored in teal and points corresponding to Cluster 1B (positive PC 1 value) colored in sea green. (B) Per-residue RMSF plot of all Cluster 1 protomers. Significant increase between A85-F94 relative to rest of sequence determines subcluster membership. Representative Cluster 1A protomer 2N0A (C) and Cluster 1B protomer 6A6B (D) shown with I88 and F94 sidechains rendered in black.

Cluster 1A contains all four motifs described above and illustrated in **Figure 3**: (1) an intermolecular salt-bridge forming between E46-K80 (**Figure 3F**), (2) a glutamine ladder at Q79 (**Figure 3E**), (3) steric zippers forming at A78-V49-A76, A53-V74 (**Figure 3C**) and V77-A90-T75-T92-V71 (**Figure 3D**), and (4) a closed-in hydrophobic pocket placing the sidechains of I88, A91 and F94 in close proximity, producing a tight fold with an average RMSF of 1.19 Å (**Figure 4C and S3A**). Cluster 1B also contains (1) an intermolecular salt-bridge forming between E46-K80, (2) a glutamine ladder at Q79, (3) and a steric zipper forming between A78-V49-A76-V52, however it undergoes a slight register change resulting in the formation of a C-terminal steric zipper between V74-A89-V71-A91-A69, opening up the hydrophobic pocket and disrupting the close interaction of I88-A91-F94 seen in Cluster 1A (**Figure 4D and S3B**). This is present in the first Cluster 1B fibril determined by Cryo-EM (6A6B) (**Figure 4D**). Like many other Cluster 1B fibrils, 6A6B exists as a two protofilament fibril, showcasing an additional interfilament steric zipper between V551-A53_2_-A53_1_-V55_2_. While in all Cluster 1B fibrils, the sidechain of I88 does not interact with F94, it does interact with the sidechain of V77 in all cases except for 6OSM, where it is exposed to the solvent. Additionally, the F94 sidechain is solvent exposed in all structures but is close in proximity to G67 (and on the same side of the beta strand as I88) in all cases except for the *ex vivo* MSA fibrils (6XYO_C, 6XYP_C and 6XYQ_C), the *in vitro* JOS-like MAAAEKT insertion fibrils (8CE7 and 8CEB) and the *ex vivo* seeded PD CSF fibrils (7V49 and 8H05) **(Figure S3B)**. The lack of I88-A91-F94 interaction in Cluster 1B structures constitutes a decrease in tight sidechain packing and conservation, substantiated by the decreased density of the Cluster 1B data points in PC space as compared to those of Cluster 1A and an average RMSF of 2.14 Å **(Figure 4A-B)**. Interestingly, the *ex vivo* MSA fibrils (6XYO, 6XYP and 6XYQ) each have one filament from Cluster 1A and one filament from Cluster 1B, suggesting that a change in the steric zipper register and I88-A91-F94 interaction has little effect on the rest of the fibril structure. This pocket may serve as a template for targeting by therapeutic ligands (**Figure S6**).

### CLUSTER 2 STRUCTURES

Structures of Cluster 2 fibrils have exclusively been determined by Cryo-EM and include 12 *in vitro* fibril preparations, six *ex vivo* seeded MSA structures, and nine *ex vivo* seeded PD structures. The structures have an average RMSF of 1.99 Å (**Figure S2D**) and are illustrated in **Figure 5A** as an alignment with 6RT0 (the first determined structure of Cluster 2). The RMSF only increases above 3.0 Å for each residue between G84-A91 (**Figure S2B**). In the same framework of stability contributions, we applied to Cluster 1 fibrils (**Figure 3**), all Cluster 2 fibrils contain (1) a glutamine ladder at Q79 (**Figure 5E**) and (2) a steric zipper between T72-V49-V74 (**Figure 5D**). They do not contain a Greek key motif spanning A69-F94.

**Figure 5:**
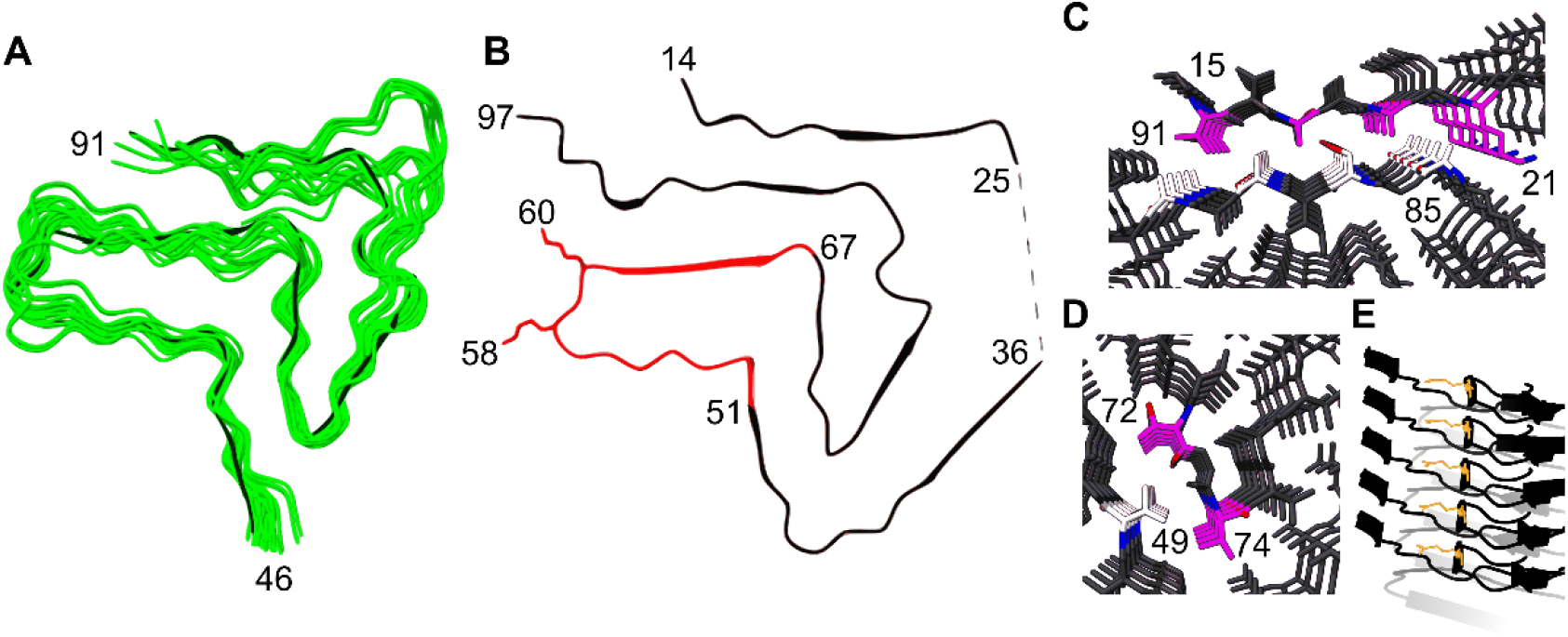
Characteristic features of Cluster 2 polymorphs. (A) Overlay of all Cluster 2 structures in green with 6RT0 in black, aligned from E46-A91. (B) Full length peptide of 6RT0 showing G51-K58-K60-G67 β-arc motif in red with protruded K58 and K60 sidechains pointing out towards the solvent. Steric zippers forming between (C) A91-V15-A89-A17-S87-A19-A85-K21 and (D) T72-V49-V74, with carbons on opposing beta strands colored in purple and white, nitrogen atoms colored in blue, and oxygen atoms colored in red. (E) Side view of the Q79 ladder (orange).

Instead, they feature a prominent, conserved, β-arc motif between G51-G67 marked by the sidechains of K58 and K60 pointing away from the fibril core exposed to the solvent (**Figure 5B**). In addition, 11 of the 27 Cluster 2 fibrils (8A4L, 8ADS, 8AEX, 7V4A, 7V4B, 7OZG, 7NCI, 7NCH, 6SSX, 6SST, 6RT0) contain an ordered β-strand in the N-terminus, from V15-K21, which forms a steric zipper between A91-V15-A89-A17-S87-A19-A85-K21 (**Figure 5C**).

We repeated the subcluster analysis on the protomers in Cluster 2 (**Figure 6A**), which revealed three subclasses, each of which have slight differences in the angle at G84 and the corresponding register of the G84-A91 stretch. The per-residue RMSF is highest between G84-A91, reflecting these inter-subclass differences, but also slightly increases at G67 relative to its nearby residues (**Figure 6B**). Cluster 2A, illustrated using 6UFR (**Figure 6C**) and as an overlay in **Figure S3C**, is highly conserved, with an average RMSF value of 0.67 Å. The sidechains of I88 and N65 do not interact and are noticeably far apart from one another in the structure. All Cluster 2A fibrils have an extended fibril core (ending with K97 at the C-terminus) and form a salt bridge between E61-K96 (**Figure S4A**). Cluster 2B, illustrated using 6RT0 (**Figure 6D**) and as an overlay in **Figure S3D**, is slightly less conserved than Cluster 2A, with an average RMSF of 0.77 Å. The kink in Cluster 2B structures at T81 enables the formation of a salt bridge between K80-E83 (**Figure S4B**) which, in turn, flips the A85, S87, A89 and A91 sidechains to the opposite side of their beta strand, allowing them to form a steric zipper with V15, A17, A19, and K21 on the N-terminal beta strand in fibrils where density is modeled in that region (**Figure 5C**). This results in an interaction between the I88 and N65 sidechains, changing the local conformation and increasing the local per-residue RMSF among all Cluster 2 fibrils (**Figure 6B**). Cluster 2C, illustrated using 8A4L (**Figure 6E**) and as an overlay in **Figure S3E**, is the least conserved out of the Cluster 2 subclasses, with an average RMSF of 1.15 Å. These fibrils also contain a kink at T81 and corresponding K80-E83 salt bridges (similar to Cluster 2B), but do not have an interaction between the sidechains of N65 and I88 (**Figure S4C**). The sidechain orientations are well conserved in all of Cluster 2C fibrils except for 8AEX, where the I88 sidechain is exposed to the solvent. Many of the Cluster 2 protomers exist in two and three protofilament assemblies, which could be biased due to buffer conditions or cofactors, but do not sort cleanly by subclass. It is notable that the average RMSF values for each subclass are significantly lower than any of the EM-map resolutions of their member structures (**Table S1 and Figure S3C-E**), given that many of the structures were determined from independent fibril preparations by different research groups.

**Figure 6:**
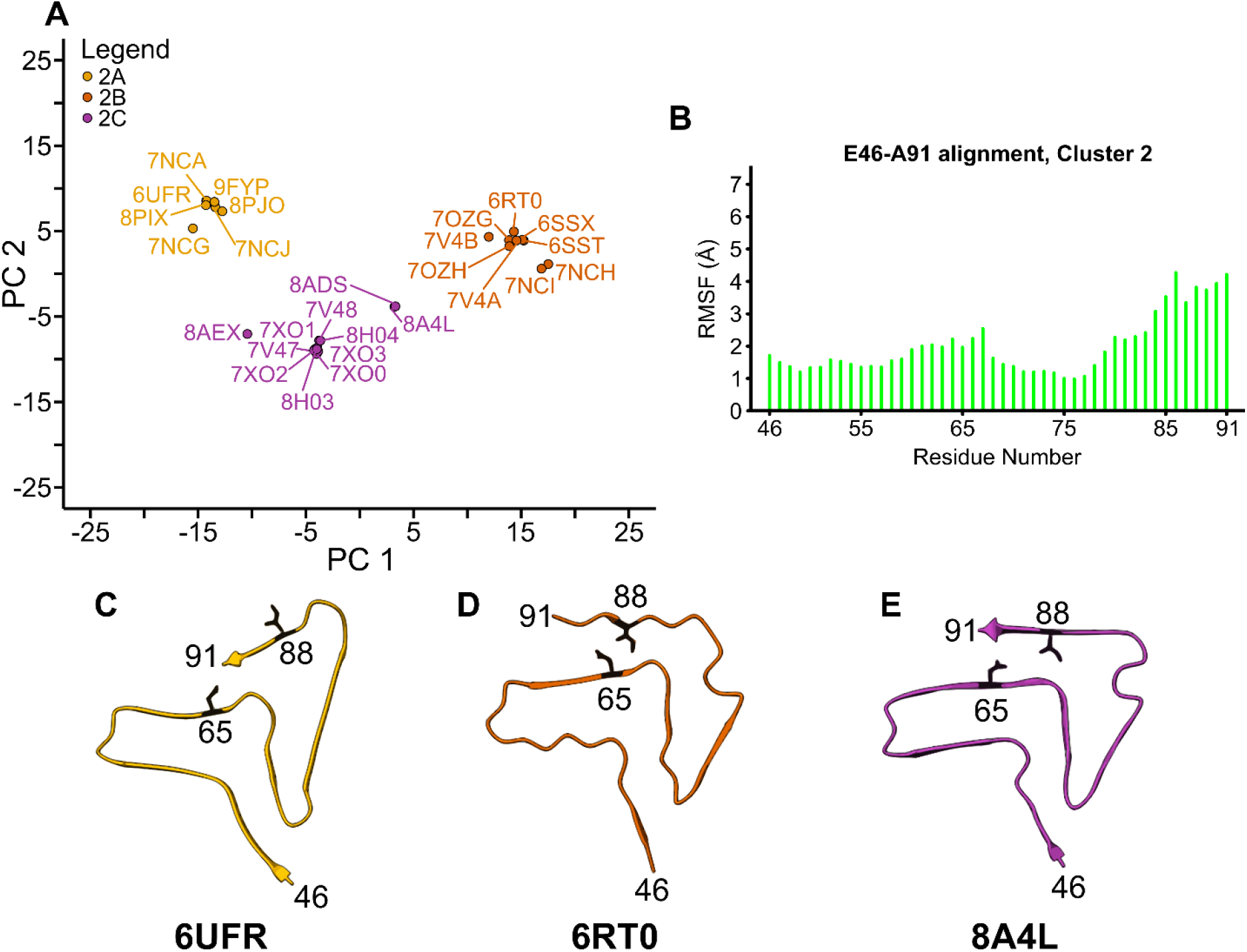
Identification of subclasses within Asyn fibril Cluster 2 protomers. (A) PCA plot (PC 2 vs PC 1) of all protomers within Cluster 2. Each point represents the protomer of single PDB coordinate file and is labeled with its four-character identifier, with points corresponding to Cluster 2A, Cluster 2B and Cluster 2C colored in orange, red and magenta, respectively. (B) RMSF vs. residue number is plotted for all Cluster 2 protomers, showing a significant increase in RMSF between A85-A91 and a slight local increase at G67. (C) Representative Cluster 2A protomer (6UFR), (D) Cluster 2B protomer (6RT0) and (E) Cluster 2C protomer (8A4L) are shown with the N65 and I88 sidechains rendered in black, showing the residues and beta-strands that differ the most between subclasses.

### UNCLASSIFIED STRUCTURES CONTAIN CONSERVED MOTIFS

The unclassified structures include 11 *in vitro* preparations, one *ex vivo* LBD structure and one *ex vivo* seeded LBD structure. The structures have an average RMSF of 8.61 Å (**Figure S2F**) and are illustrated as an overlay in **Figure S2G**. Many of these structures may appear close in PC space due to identical preparation methods, including those with residue Y39 phosphorylated (6L1U and 6L1T) (51) and those formed in the presence of lipid vesicles (8ADV, 8ADU and 8ADW) (28). Those with familial mutation E46K and E46K cross seeded with WT (6L4S and 7C1D, respectively) (50,54) also appear close in PC space (**Figure 2D**). Fibrils formed under independent aggregations but with identical constructs and conditions (8PK2 and 8PK4) (31) strikingly formed different structures, such that 8PK2 is found in Cluster 1A while 8PK4 remains unclustered and close in PC space to the pY39 fibrils. Other structures appear further away from the bulk of unclustered structures (PC 1 > -30), with 7LC9_C (55) and 8GF7 (61) situated nearby but outside of the PC space encompassing Cluster 1 fibrils. The 8GF7 fibrils were formed with the S87 O-GlcNac post-translational modification, which introduces a bulky sidechain, remodeling the Greek key region and affecting the structure of the entire N-terminus as well. The 7LC9 fibrils have two asymmetric protofilaments, one denoted by 7LC9_F (sequence did not span E46-A91 so was not initially clustered (**Figure S1**)) and one by 7LC9_C. The 7LC9_C protomer has some similarities in structure to Cluster 1 fibrils, particularly sharing the same turn at G67-G68, but mostly disagrees, with an average RMSF > 5 Å when compared to representative structures. One of the fibrils formed in the presence of heparin (7V4C) (27) is situated directly in between Clusters 1 and 2 along the PC2 axis (**Figure 2A**). This fibril contains both a Greek key motif similar to that in Cluster 1B (all-atom RMSD of 2.96 Å to 6OSM (48) for residues 69-91) and a β-arc motif consistent with Cluster 2 fibrils (all-atom RMSD of 1.40 Å to 6SSX (47) for residues 51-67). The two unclustered point mutation structures, E46K (6L4S) (50) and WT cross-seeded with E46K (7C1D) (54) are also situated directly in between Clusters 1 and 2 along the PC2 axis. Both of these structures contain Greek key motifs similar to Cluster 1A fibrils (all-atom RMSD of 2.36 Å (6L4S) and 1.61 Å (7C1D) to 2N0A (32) for residues 69-91) and a β-arc motif consistent with Cluster 2 fibrils (all-atom RMSD of 1.50 Å (6L4S) and 1.82 Å (7C1D) to 6SSX (47) for residues 51-67).

Due to the existence of conserved motifs in some of the unclassified structures, we re-clustered the structures based on subsets of their residues – G51-G67 (**Figure S5A**) and A69-A91 (**Figure S5D**). This allowed us to include previously unaligned structures which did not have fibril cores spanning E46-A91, including 6CU8 (43), 8JEX (63), 8JEY (63), 6RTB (47), 8RI9 (62), and 7LC9_F (55). By using a similar approach as before, we aligned the structures, reduced their dimensionality using PCA and identified clusters within the G51-G67 structures. This revealed 2 major clusters, one of which forms a backwards C-shaped structure (previously referred to as part of a *boot* motif (37)) with a turn at T59 (Cluster 1) and the other formed the characteristic β-arc from Cluster 2 (**Figure S5A**). The Cluster 1 structures include all Cluster 1 structures from **Figure 2** (average RMSF of 1.29 Å) and the Cluster 2 structures include all the Cluster 2 structures from **Figure 2** (average RMSF of 0.88 Å). A majority of the previously unaligned and unclassified structures became part of a cluster here; specifically, 6CU8 and 7LC9_C joined Cluster 1 (**Figure S5B**) and 8ADU, 8ADV, 8ADW, 6L4S, 7E0F, 7V4C, 7C1D, 8A9L, 6L1T, 6L1U, 8JEX and 8PK4 joined Cluster 2 (**Figure S5C**). Only three out of the 72 structures in this alignment remained unclassified, suggesting the high degree of structural conservation within this G51-G67 region of Asyn fibril cores. For one of the unclassified structures, 8FPT_1, the region between G51-G67 was found to be dynamically disordered and not amenable to residue-specific assignment and confident structure determination within the SSNMR data. Recent findings detailing the structure of Asyn oligomers formed in the presence of phospholipid vesicles suggest that the β-arc between G51-G67 is was formed during the early events of fibril assembly, providing a structural basis for this conserved conformation across independent fibril preparations (75).

We next applied the same approach and clustered the A69-A91 structures, which cleanly separated the fibrils by subclass, as expected from the RMSF analyses (**Figure S5D**). Previously unaligned and unclassified structures were able to join each subclass, with 7C1D, 6L4S, 7E0F and 8GF7 joining Cluster 1A (1.24 Å average RMSF) (**Figure S5E**), 8RI9 and 7V4C joining Cluster 1B (1.81 Å average RMSF) (**Figure S5F**) and 6RTB (previously not clustered due to missing residues between A56-E62) (47) joining Cluster 2B/C (1.47 Å average RMSF) (**Figure S5H**). While 8GF7 joins Cluster 1A, it differs from the other 15 structures in that the S87 O-GlcNac modification positions the S87 sidechain in contact with F94 and flips the I88 sidechain in towards the remainder of the fibril core. That several other structures join Clusters 1A and 1B here, each containing a Greek key motif, suggests the high degree of structural conservation in the A69-A91 region of the fibril core. A previous time-resolved structural study of Asyn fibril formation in the presence of phospholipid vesicles suggests that residues in this Greek key region are the first to form β-sheets (12), providing an experimental rationalization for this conserved conformation across independent fibril preparations.

The only structure that does not contain any conserved motifs is 8JEY (phosphorylated S87) (63). The fibril core of this structure spans M1-T72, excluding S87 and much of the hydrophobic region which remain part of the fibril’s mobile and heterogeneous fuzzy coat. It is essential to note that the majority of the fibrils used in this study were untwisted fibrils, yet helical reconstruction was applied to the minority population of *twisted* fibrils (∼26%). It is possible – as the reviewers of that study have noted (see peer review file (63)) – that the untwisted fibril majority contain motifs found in wild-type structures. Indeed, it is alternatively possible that the introduction of a significant negative charge within the fibril core can induce atypical fibril formation.

### OUTLOOK ON LEWY FOLD STRUCTURES

For the A69-A91 alignment, 7LC9_F (41-140 truncation), 8A9L (*ex vivo* LBD) and 8FPT_1 (*ex vivo* seeded LBD) are close together in PC space (**Figure S5D**) and have a high structural agreement with one another (1.26 Å average RMSF) (34,55,58). The entire structure of 8A9L has been referred to as the Lewy fold (34), but we see this strong agreement only when analyzing the subset of residues and not the entire fibril. Even though the structures are close in PC space to Cluster 1A, their I88 and F94 sidechains do not interact and the turn at G73 is in the opposite direction, 180° apart (**Figure S7**). There is a turn at E83, suggesting that residues G73-A91 form an extended β-arc. Because there are only three of these Lewy fold-type fibrils and our clustering algorithm would overfit the data if the minimum cluster size was set at three (see supporting material), even though they all contain a conserved β-arc, they do not form a cluster. Indeed, it is significant that this G73-A91 β-arc motif is found in one *in vitro* preparation and two patient-derived (*ex vivo* and *ex vivo* seeded) LBD fibrils, suggesting structure-disease specificity. Even though this motif was only replicated *in vitro* in one specific study with N-terminally truncated Asyn (55), it suggests that it is a kinetically accessible motif somewhere on the *in vitro* fibril formation trajectory. It also demonstrates the importance of recapitulating structures containing this motif *in vitro* to develop reproducible and scalable models for studying LBD pathogenesis and testing therapeutic ligands.

## CONCLUSION

We have shown that aligning and clustering the published Asyn protomers by their fibril cores classifies Asyn tertiary structure into two major classes, accounting for 81% of the structures. Cluster 1 is characterized by a Greek key motif spanning residues 69-95, and Cluster 2 is characterized by a β-arc motif spanning residues 51-67. These two classes have intraclass variations which are largely defined by structural heterogeneity within the stretch of residues between A85-A91. Cluster 1 subclasses are distinguished based on the interaction (or lack thereof) of the I88, A91 and F94 sidechains. The *ex vivo* JOS fibrils (8BQV and 8BQW) display such an interaction and therefore fall into Cluster 1A, and the *ex vivo* MSA fibrils (6XYO, 6XYP and 6XYQ) each have two asymmetric filaments, one of which falls into Cluster 1A and the other which falls into Cluster 1B (33,59). Cluster 2 subclasses are distinguished based on both the formation of a steric zipper between the A85-A91 beta strand and the V15-K21 beta strand (Cluster 2B) as well as salt bridge formation either between E61-K96 (Cluster 2A) or between K80-E83 (Cluster 2B and 2C).

As we have shown, 16 *in vitro* preparations of Cluster 1A and 1B fibrils have been well documented and published, and while the exact molecular determinants which bias the formation of each subcluster are still unknown, Cluster 1A and 1B fibrils make attractive *in vitro* models of MSA. A recent preprint has shown that fibrils of such type can reproduce MSA-like inclusions in mouse brains (76), suggesting further utility as models for MSA. The overall process for recombinant expression of *in vitro* Asyn monomer and growth of fibrils, resulting in structures of Cluster 1A and 1B, is well established and does not require difficult-to-obtain diseased tissue samples. *In vitro* preparation also yields fibrils on the milligram scale rather than the picogram scale, which is crucial if Asyn fibrils are to be used as substrates for drug development and in uncovering the molecular mechanisms behind fibril formation in disease.

Lastly, by re-clustering the structures based on sequence subsets where conserved motifs are present, we showed that all but one structure (8JEY) contains a conserved Greek key or β-arc motif. Along with experimental data from SSNMR studies, this finding suggests that these motifs are likely formed during the early events of fibril assembly. As such, designing ligands and diagnostic tools which specifically bind to these structural motifs can help identify *in vivo* fibril formation mechanisms and target disease-specific fibrils in their nascent forms.

## ACKNOWLEDEMENTS

This work was supported by 2RF1NS110436-06 (NINDS), P41GM136463 (NIGMS), the William H. Peterson Graduate Fellowship in Biochemistry (awarded to M.H.M.), the Molecular Biophysics Predoctoral Training Grant T32GM130550 (NIGMS) (awarded to O.A.W.), and the NIH Ruth L. Kirschstein Fellowship F32GM149118 (NIGMS) (awarded to C.G.B.). We would like to thank Dr. Juan Sanchez, Dr. Joshua Pierson, Benjamin Harding, Prof. Hannah Wayment-Steele, Dr. Songlin Wang and Spencer Hyman for their helpful discussions and suggestions.

