## Supporting Material for "Alpha-Synuclein Fibril Structures Cluster into Distinct Classes"

**SUPPLEMENTAL METHODS**

**PDB alignment**

Protein Data Bank (PDB) files were downloaded from the Research Collaboratory for Structural Bioinformatics (RCSB) and edited within ChimeraX to remove protons and were saved as individual protomers (1,2). The Bio3D package in the R programming language was used to import and align the protomer PDB files using MUltiple Sequence Comparison by Log-Expectation (MUSCLE) to make their CA-coordinates invariant to one another (3,4). We used a subset of the initial protomers (68 total) in our alignment due to missing residues in some protomers (**Figure S1**). Residues E46-A91 were aligned for all Principal Component Analysis (PCA) figures in the main text. Residues G51-G67 and residues A69-A91 were aligned for PCA in Figure S5A and S5D, respectively.

**Dimensionality reduction using PCA**

We used PCA to reduce the dimensions of our PDB coordinates down to two so that we could easily visualize their similarities and differences on an x-y plane (3,5). This was specifically done for the alignment of protomers between residues E46-A91 for **Figure 2** and **Figure 6**, residues E46-F94 for **Figure 4**, residues G51-G67 for **Figure S5A** and residues A69-A91 for **Figure S5D**. PCA can be implemented within the R programming language quite easily and integrates nicely with the Bio3D formatted data.

**DBSCAN clustering**

We used Density-Based Spatial Clustering of Applications with Noise (DBSCAN) to cluster protomers based on their coordinates (6). DBSCAN clustering is available as a function within the R programming language (7), and the algorithm requires the following two values as input from the user: *minPts*, the minimum number of points in the *epsilon*-neighborhood of the cluster, and *epsilon,* the radius from a core point which reaches other points within the cluster. *minPts* was chosen to be 6, following the method of Sander et al. such that *minPts* = 2*nDim (8). We chose *epsilon* by following the method of Ester and Kriegel in their original DBSCAN paper to compute the k-nearest neighbor distances of each protomer and plot that vs. the number of protomers and find the point of largest curvature, or *elbow,* in the plot. At that point in x (number of protomers), the k-nearest neighbor distance should be the value of *epsilon* chosen for DBSCAN.

**SUPPLEMENTAL FIGURES AND TABLES**

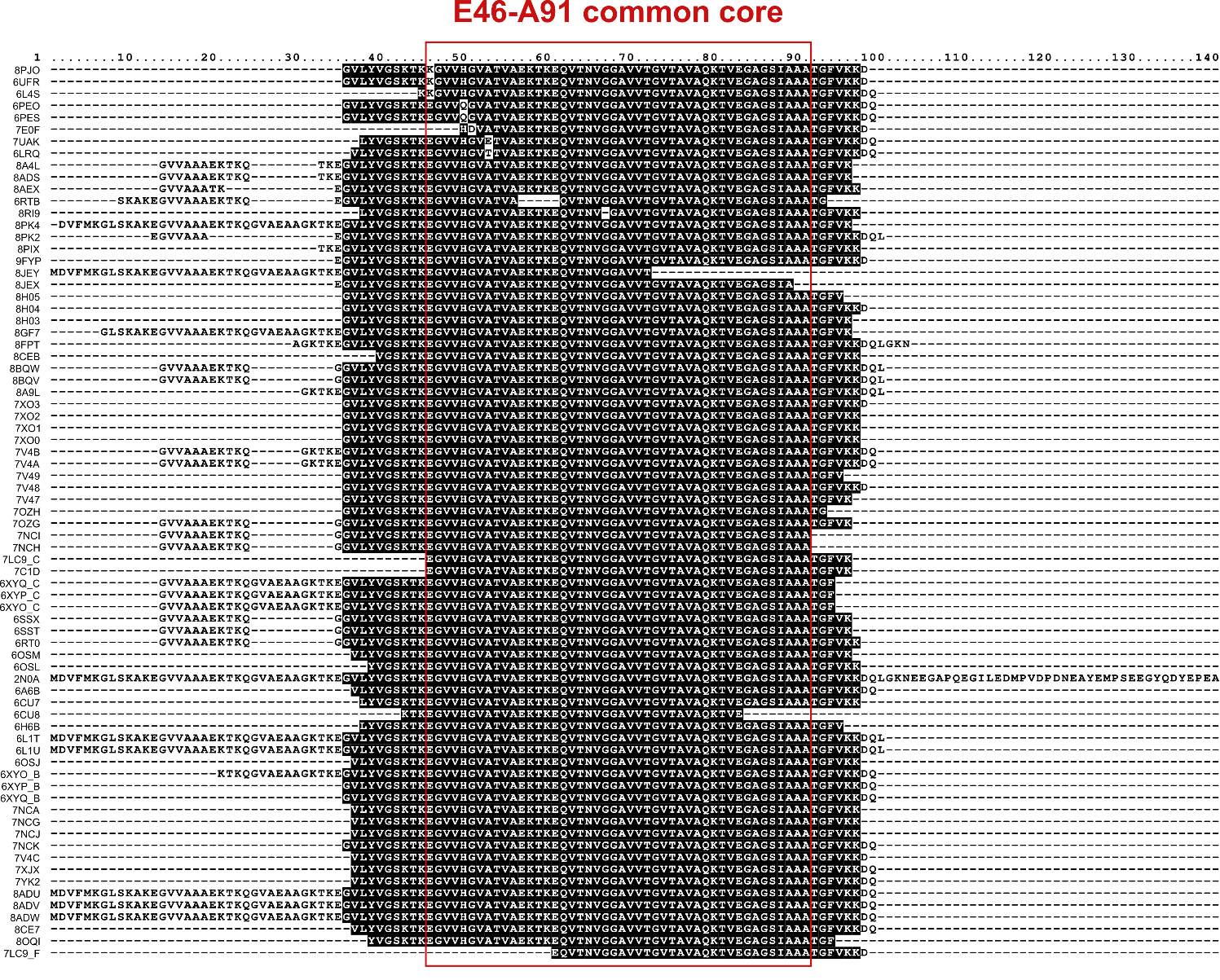

**Figure S1:** Multiple Sequence Alignment of Asyn protomers. Each protomer is listed with its modeled sequence listed out between M1-A140. Blacked out residues span from G36-K97, which form the fibril core in structures which contain those modeled residues. Unshaded residues within the fibril core of particular PDBs (8PJO, 6UFR, 6L4S, 6PEO, 6PES, 7E0F, 7UAK, and 6LRQ) indicate hereditary mutations. The region used for DBSCAN clustering (**Figure 2**) is boxed in red and indicated as E46-A91. PDBs which don’t have residues between E46-A91 modeled were excluded from the clustering analysis in the main text but included for the analysis in **Figure S5**.

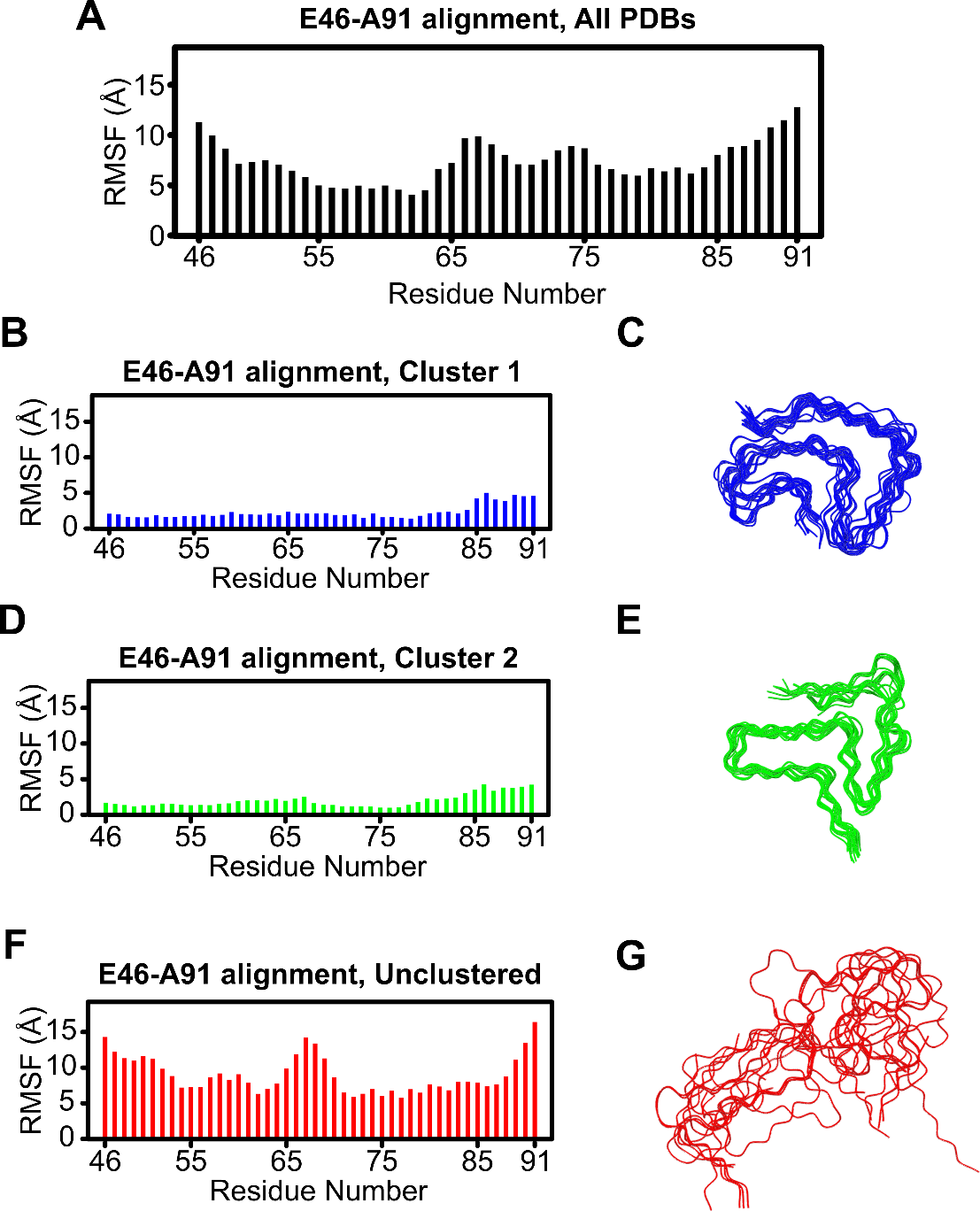

**Figure S2:** Per-residue root-mean-square deviation (RMSF). (A) Per-residue RMSF for all structures. (B) Cluster 1 per-residue RMSF (2.31 Å average) and (C) protomer alignment. (D) Cluster 2 per-residue RMSF (1.99 Å average) and (E) protomer alignment. (F) Unclassified protomers per-residue RMSF (8.61 Å average) and (G) protomer alignment.

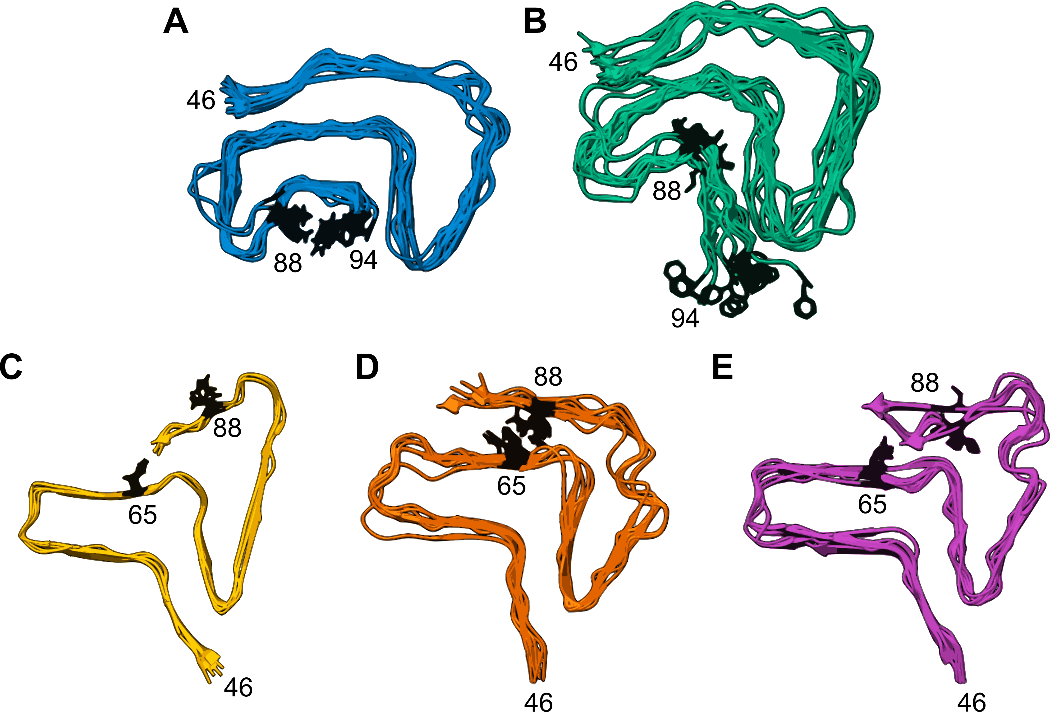

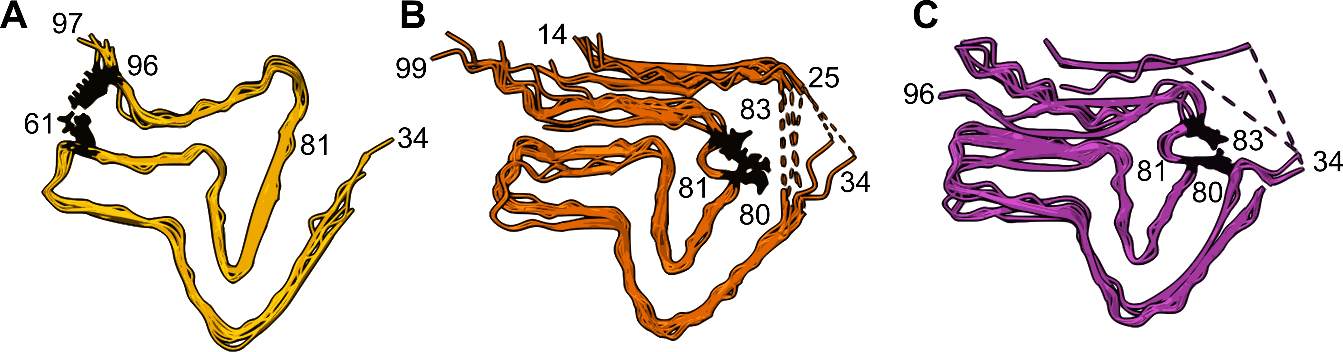

**Figure S3:** Overlays of fibril subclasses with key sidechains (I88 and F94 for Cluster 1, N65 and I88 for Cluster 2) rendered in black. (A) Subclass 1A has an average RMSF of 1.19 Å, (B) Subclass 1B has an average RMSF of 2.14 Å, (C) Subclass 2A has an average RMSF of 0.67 Å, (D) Subclass 2B has an average RMSF of 0.77 Å, and (E) Subclass 2C has an average RMSF of 1.15 Å.

**Figure S4:** Salt bridge formation in Cluster 2 fibrils. (A) Cluster 2A fibrils overlaid with the E61-K96 salt bridge rendered and colored in black. (B) Cluster 2B fibrils overlaid with the K80-E83 salt bridge colored in black. (C) Cluster 2C fibrils overlaid with the K80-E83 salt bridge colored in black.

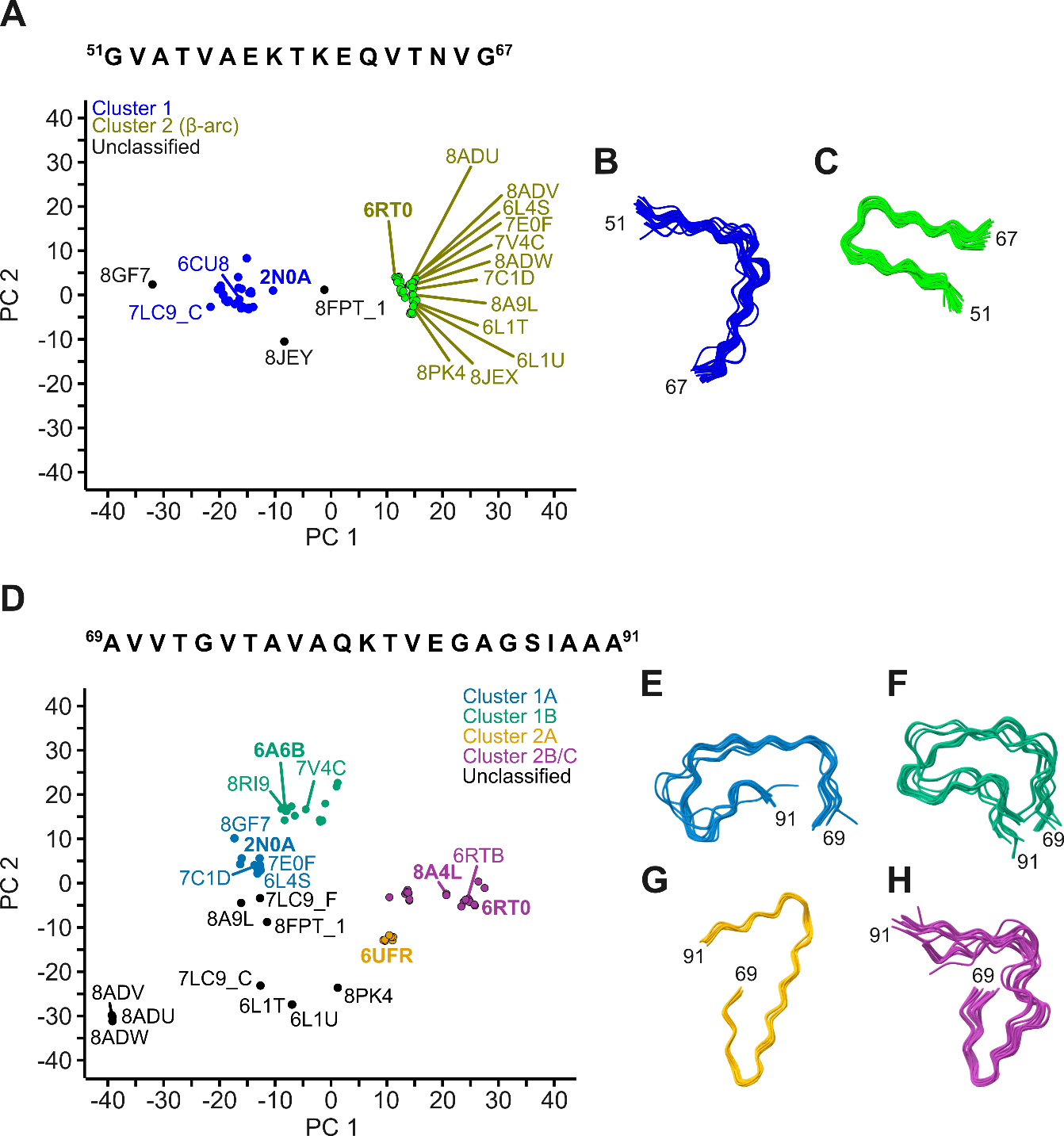

**Figure S5:** Clustering based on conserved sequence motifs. (A) Alignment and PCA of all structures in Table S1 which have sequences between G51-G67. DBSCAN used to cluster structures, with blue points denoting membership in Cluster 1, green points for Cluster 2, and black points as unclassified. 2N0A and 6RT0 are bolded as they were the first-experimentally determined structures in their respective clusters. Labeled points correspond to previously unclassified structures, where the majority now fall into a cluster. (B) Protomer overlay of Cluster 1 structures. (C) Protomer overlay of Cluster 2 structures, detailing the conserved β-arc motif. (D) Alignment and PCA of all structures in Table S1 which have sequences between A69-A91. DBSCAN used to cluster structures, with Cluster 1A colored in teal, Cluster 1B colored in sea green, Cluster 2A colored in orange, Clusters 2B/C colored in magenta, and the unclassified structures colored in black. (E-H) Protomer alignments of each cluster, with Greek key motifs shown in (E) and (F).

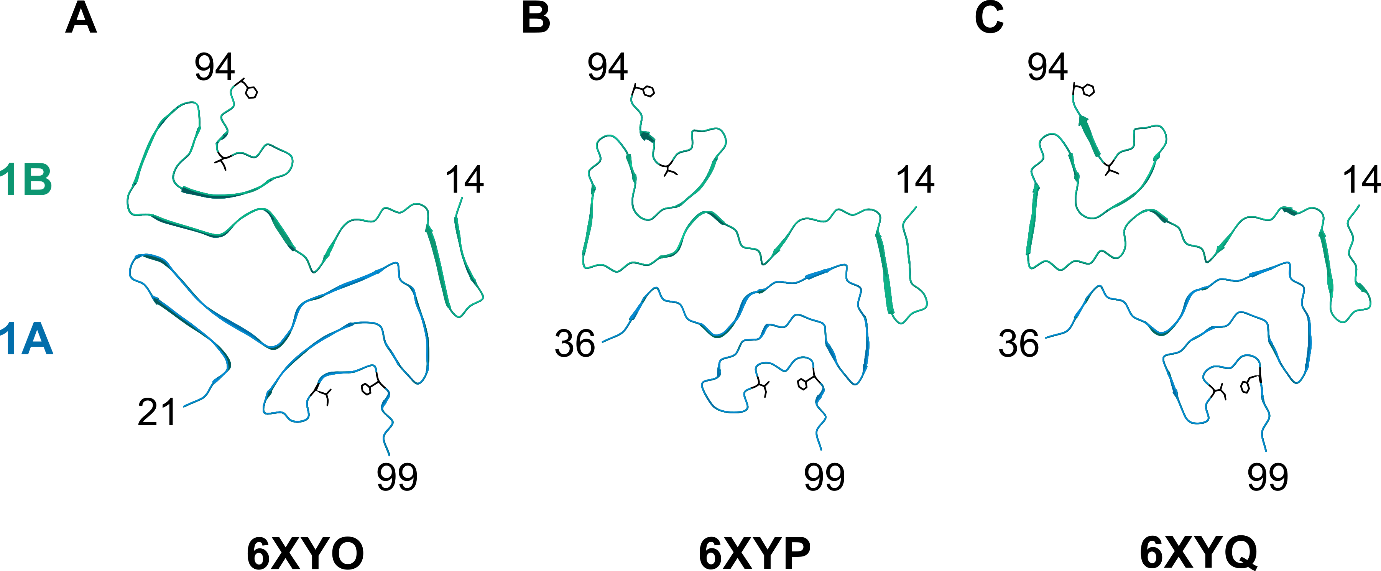

**Figure S6:** Structures of ex vivo MSA fibrils. (A) 6XYO, (B) 6XYP and (C) 6XYQ are shown as 2 protofilaments each. For each structure, one protofilament is classified in Cluster 1A (teal and the other is classified in Cluster 1B (sea green). The sidechains of I88 and F94 are rendered and colored in black, both pointing out of the fibril core and interacting with each other in the Cluster 1A protomer and pointing in opposite directions, not interacting, in the Cluster 1B protomer.

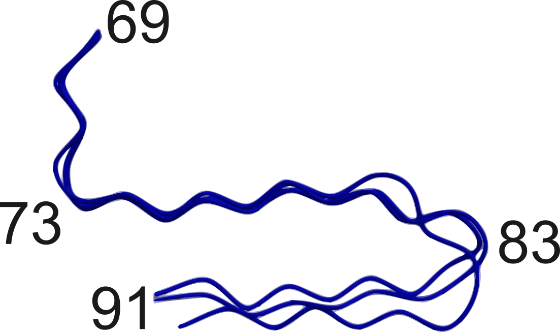

**Figure S7:** Alignment of Lewy Fold structures, from A69-A91. Structures 8FPT_1, 8A9L and 7LC9_F are aligned and colored in navy, with select residues labeled.

| PDB Code | Protomer Chain ID | Preparation Method | Construct Details | Determination Method | Resolution (Å) | Reference |
| --- | --- | --- | --- | --- | --- | --- |
| **2N0A** | C | *In vitro* | WT | SSNMR | N/A | (9) |
| **6PEO** | C | *In vitro* | H50Q | Cryo-EM | 3.30 | (10) |
| **6PES** | C | *In vitro* | H50Q | Cryo-EM | 3.60 | (10) |
| **6LRQ** | C | *In vitro* | A53T | Cryo-EM | 3.49 | (11) |
| **6L4S** | C | *In vitro* | E46K | Cryo-EM | 3.37 | (12) |
| **7NCK** | C | *Ex vivo* seeded | MSA | Cryo-EM | 3.18 | (13) |
| **7C1D** | C | *In vitro* | WT/E46K | Cryo-EM | 3.80 | (14) |
| **7E0F** | C | *In vitro* | G51D | Cryo-EM | 3.02 | (15) |
| **8BQV** | A | *Ex vivo* | JOS | Cryo-EM | 2.00 | (16) |
| **8BQW** | C | *Ex vivo* | JOS | Cryo-EM | 2.30 | (16) |
| **7UAK** | C | *In vitro* | A53E | Cryo-EM | 3.38 | (17) |
| **8PK2** | C | *In vitro* | WT | Cryo-EM | 3.26 | (18) |
| **6XYQ** | B | *Ex vivo* | MSA | Cryo-EM | 3.09 | (19) |
| **6XYO** | B | *Ex vivo* | MSA | Cryo-EM | 2.60 | (19) |
| **6XYO** | C | *Ex vivo* | MSA | Cryo-EM | 2.60 | (19) |
| **6XYP** | C | *Ex vivo* | MSA | Cryo-EM | 3.29 | (19) |
| **6XYQ** | C | *Ex vivo* | MSA | Cryo-EM | 3.09 | (19) |
| **6XYP** | B | *Ex vivo* | MSA | Cryo-EM | 3.29 | (19) |
| **6CU8** | C | *In vitro* | WT | Cryo-EM | 3.60 | (20) |
| **8GF7** | C | *In vitro* | S87 O-GlcNAc | Cryo-EM | 4.80 | (21) |
| **6OSM** | C | *In vitro* | 1-103 | Cryo-EM | 3.40 | (22) |
| **7V49** | C | *Ex vivo* seeded | PD | Cryo-EM | 3.40 | (23) |
| **8H05** | C | *Ex vivo* seeded | PD | Cryo-EM | 3.40 | (23) |
| **7V4C** | F | *In vitro* | WT | Cryo-EM | 3.30 | (24) |
| **6A6B** | C | *In vitro* | WT | Cryo-EM | 3.07 | (25) |
| **6H6B** | C | *In vitro* | WT | Cryo-EM | 3.40 | (26) |
| **6CU7** | C | *In vitro* | WT | Cryo-EM | 3.50 | (20) |
| **6OSJ** | C | *In vitro* | WT | Cryo-EM | 2.80 | (22) |
| **6OSL** | C | *In vitro* | 1-122 | Cryo-EM | 3.00 | (22) |
| **8RI9** | C | *In vitro* | WT | Cryo-EM | 3.30 | (27) |
| **7XJX** | C | *In vitro* | WT | Cryo-EM | 2.70 | (28) |
| **8OQI** | C | *In vitro* | WT | Cryo-EM | 3.10 | (29) |
| **8CE7** | C | *In vitro* | 22-MAAAEKT insertion | Cryo-EM | 2.70 | (16) |
| **8CEB** | C | *In vitro* | 22-MAAAEKT insertion | Cryo-EM | 2.80 | (16) |
| **7YK2** | C | *In vitro* | WT | Cryo-EM | 2.80 | (30) |
| **6SSX** | C | *In vitro* | WT | Cryo-EM | 2.98 | (31) |
| **6RT0** | C | *In vitro* | WT | Cryo-EM | 3.1 | (31) |
| **6RTB** | C | *In vitro* | WT | Cryo-EM | 3.46 | (31) |
| **6SST** | C | *In vitro* | WT | Cryo-EM | 3.40 | (31) |
| **7NCH** | C | *Ex vivo* seeded | MSA | Cryo-EM | 3.84 | (13) |
| **7NCI** | C | *Ex vivo* seeded | MSA | Cryo-EM | 3.55 | (13) |
| **7OZG** | C | *Ex vivo* seeded | PD | Cryo-EM | 3.30 | (32) |
| **7OZH** | C | *Ex vivo* seeded | MSA | Cryo-EM | 3.02 | (32) |
| **7V47** | C | *Ex vivo* seeded | PD | Cryo-EM | 2.80 | (23) |
| **7V48** | C | *Ex vivo* seeded | PD | Cryo-EM | 3.00 | (23) |
| **7V4A** | C | *In vitro* | WT | Cryo-EM | 3.20 | (24) |
| **7V4B** | C | *In vitro* | WT | Cryo-EM | 3.10 | (24) |
| **8A4L** | C | *In vitro* | WT | Cryo-EM | 2.68 | (33) |
| **8ADS** | C | *In vitro* | WT | Cryo-EM | 3.05 | (33) |
| **8AEX** | C | *In vitro* | WT | Cryo-EM | 2.76 | (33) |
| **7XO0** | C | *Ex vivo* seeded | PD | Cryo-EM | 3.00 | (23) |
| **7XO1** | C | *Ex vivo* seeded | PD | Cryo-EM | 3.00 | (23) |
| **7XO2** | C | *Ex vivo* seeded | PD | Cryo-EM | 3.00 | (23) |
| **7XO3** | C | *Ex vivo* seeded | PD | Cryo-EM | 2.60 | (23) |
| **8H03** | C | *Ex vivo* seeded | PD | Cryo-EM | 2.80 | (23) |
| **8H04** | C | *Ex vivo* seeded | PD | Cryo-EM | 3.00 | (23) |
| **6UFR** | C | *In vitro* | E46K | Cryo-EM | 2.50 | (34) |
| **9FYP** | C | *In vitro* | WT | Cryo-EM | 2.23 | (18) |
| **8PIX** | C | *In vitro* | WT | Cryo-EM | 3.41 | (18) |
| **8PJO** | C | *In vitro* | E46K | Cryo-EM | 2.31 | (18) |
| **7NCA** | C | *Ex vivo* seeded | MSA | Cryo-EM | 3.47 | (13) |
| **7NCG** | C | *Ex vivo* seeded | MSA | Cryo-EM | 3.43 | (13) |
| **7NCJ** | C | *Ex vivo* seeded | MSA | Cryo-EM | 4.23 | (13) |
| **8A9L** | A | *Ex vivo* | PD/DLB | Cryo-EM | 2.20 | (35) |
| **7LC9** | F | *In vitro* | 41-140 | Cryo-EM | 3.20 | (36) |
| **6L1T** | C | *In vitro* | pY39 | Cryo-EM | 3.22 | (37) |
| **6L1U** | C | *In vitro* | pY39 | Cryo-EM | 3.37 | (37) |
| **8PK4** | C | *In vitro* | WT | Cryo-EM | 3.30 | (18) |
| **8FPT** | C | *Ex vivo* seeded | DLB | SSNMR | N/A | (38) |
| **8ADU** | C | *In vitro* | WT | Cryo-EM | 3.24 | (33) |
| **8ADV** | C | *In vitro* | WT | Cryo-EM | 2.98 | (33) |
| **8ADW** | C | *In vitro* | WT | Cryo-EM | 2.95 | (33) |
| **8JEX** | B | *In vitro* | S87 O-GlcNAc | Cryo-EM | 3.10 | (39) |
| **8JEY** | C | *In vitro* | pS87 | Cryo-EM | 2.60 | (39) |
| **7LC9** | C | *In vitro* | 41-140 | Cryo-EM | 3.20 | (36) |

**Table S1:** Table of each of the 75 unique protomers published in the RCSB PDB associated with peer-reviewed articles as of June 2024. Columns reading left to right contain the 4-letter PDB identifier code, the chain name in the atomic coordinates file, the preparation method, the specifics of the construct used, determination method, and finally the resolution of the final EM map in the case of structures determined using Cryo-EM. The last column denotes the reference of the publication first reporting the PDB file.
